# Identification of genes involved in Kranz anatomy evolution of non-model grasses using unsupervised machine learning

**DOI:** 10.1101/2024.01.31.578221

**Authors:** Santiago Prochetto, Georgina Stegmayer, Anthony J. Studer, Renata Reinheimer

## Abstract

Kranz syndrome is a set of leaf anatomical and functional characteristics of species using C_4_ photosynthesis. The current model for the evolution of C_4_ photosynthesis from a C_3_ ancestor proposes a series of gradual anatomical changes followed by a biochemical adaptation of the C_4_ cycle enzymatic machinery. In this work, leaf anatomical traits from closely related C_3_, C_4_ and intermediate species (Proto-Kranz, PK) were analyzed together with gene expression data to discover potential drivers for the establishment of Kranz anatomy using unsupervised machine learning. Species-specific Self-Organizing Maps (SOM) were developed to group features (genes and phenotypic traits) into clusters (neurons) according to their expression along the leaf developmental gradient. The analysis with SOM allowed us to identify candidate genes as enablers of key anatomical traits differentiation related to the area of mesophyll (M) and bundle sheath (BS) cells, vein density, and the interface between M and BS cells. At the same time, we identified a small subset of genes that displaced together with the change in the area of the BS cell along evolution suggesting a salient role in the origin of Kranz anatomy in grasses.

**Highlight:** Kranz syndrome is a set of leaf anatomical and functional characteristics of species using C_4_ photosynthesis. We discovered, with a novel machine learning methodology, some key genes for Kranz differentiation.

## Introduction

C_4_ photosynthesis is a mechanism that has evolved in some plant species that results in the suppression of photorespiration. Photorespiration occurs during the carbon fixation process due to the dual activity of Ribulose-1,5-bisphosphate carboxylase/oxygenase (RUBISCO). To increase the concentration of CO_2_ around RUBISCO, most C_4_ species develop two specialized leaf cell types: mesophyll cells (M) and bundle-sheath cells (BS) (Hattersley, 1984; Sage *et al*., 2012). The M cells are clustered around the BS cells in a ring-like fashion, which is a hallmark of Kranz anatomy. The BS cells house RUBISCO and have unique anatomical and genetic peculiarities like the root endodermis (Slewinski *et al*., 2012). Although Kranz anatomy has long been linked to the evolution of C_4_ photosynthesis, there is an incomplete understanding of the genes involved in the differentiation of these distinct leaf anatomy.

C_4_ photosynthesis evolved independently around 70 times in angiosperms. However, most C_4_ origins are grouped into a few plant lineages suggesting that certain lineages have characteristics that facilitate the development of C_4_ photosynthesis (Sage *et al*., 2011; Christin *et al*., 2013). Phenotypic traits such as higher density of vascular bundles (VB), a higher proportion of the tissue occupied by BS cells, and duplicated genomes, have been thought of as preconditional characteristics that increase the potential to evolve to C_4_ (Sage *et al*., 2012; Sage *et al*., 2014; Christin *et al*., 2013; Christin and Osborne, 2013; Lundgren *et al*., 2019). In addition, species with C_3_-C_4_ intermediates photosynthesis are usually found near the C_4_ lineages. Indeed, current models indicate that the assembly of C_4_ photosynthesis was likely a gradual process that included the relocation of photorespiratory enzymes, and the establishment of intermediate photosynthetic subtypes. For example, species that primarily utilize the C_3_ pathway but have anatomical and physiological characteristics that reduce photorespiration, are categorized as Proto-Kranz (PK) and C_2_ type I species; while others species have an incipient C_4_ pathway, as in the C_2_ type II or the C_4_-like (Sage *et al*., 2014). The PK species have anatomical traits that tend to reduce photorespiration within BS. These include a larger area of BS, with more organelles (chloroplasts, mitochondrias, and larger vacuoles) in an arrangement similar to that observed in C_2_ species (Muhaidat *et al*., 2011). In C_2_ species, the photorespiratory pathway is partitioned between M and BS, with the release of CO_2_ occurring predominantly in BS. This occurs because of the glycine decarboxylase complex (GDC), which is responsible for the release of CO_2_ during photorespiration, and is exclusively expressed in BS mitochondria in C_2_ species (Rawsthorne *et al*., 1988). In these species, glycine formed during photorespiration in the M must diffuse or be transported to the BS to metabolize to serine, which then returns to the M to complete the photorespiratory cycle (Rawsthorne *et al*., 1988).

The grass family concentrates 60% of C_4_ species known to date which are distributed in 22-24 origins (Christin *et al*., 2013). In grasses, all species that perform C_4_ or C_2_ photosynthesis belong to the PACMAD clade. Within the PACMAD, the subtribe Otachyriinae is especially rich in different photosynthetic pathways. The subtribe is a monophyletic group consisting of 35 species divided into seven genera (Acosta *et al*., 2014, 2019). Species of the subtribe are widely distributed in humid regions of the tropics of America, Asia and Australia. In terms of photosynthetic type, the subtribe includes one C_4_ genus, and some C_2_, PK and C_3_ species (Acosta *et al*., 2014, 2019). Reconstruction of ancestral characters suggests that the common ancestor of the subtribe used the C_3_ photosynthetic pathway. The C_4_ condition would have evolved early in the diversification of the subtribe, while the extant intermediate C_2_ condition would have evolved later, prior to the diversification of the genus *Steinchisma* (Acosta *et al*., 2014). In addition, events of allopolyploidy and lateral gene transfer (LGT) that make internal classification of the subtribe difficult have been documented (Acosta *et al*., 2019). The diversity of photosynthetic types observed in the Otachyriinae subtribe make it an interesting lineage for answering multiple questions about the evolution and optimization of photosynthesis in grasses. Recently, the first genomic analysis of species from the Otachyriinae subtribe was conducted using three species of the Otachyriinae subtribe (C_3_ – *Hymenachne amplexicaulis*, PK – *Rugoloa pilosa* and C_4_ – *Anthaenantia lanata*) (Prochetto *et al*., 2023). High resolved leaf transcriptome provided in Prochetto *et al*. (2023) provides new tools for future investigation of photosynthesis evolution. Indeed, such high-quality leaf transcriptomes can be used to identify transcription factors as potential candidates for the establishment of anatomical traits related to the evolution of C_4_ photosynthesis.

For the discovery of hidden patterns of relationships between phenotypic and transcriptomic information, data integration is necessary given the need for extracting knowledge from multiple and heterogeneous data types and sources. In line with this, Self-Organizing map (SOM) tool is gaining interest among scientists by testing the potential of the method to solve multiple biological questions (Kohonen, 2013). A SOM is a special type of machine learning (ML) model, which has proven to be very well-suited for the task of heterogeneous data integration and visualization through unsupervised learning (Stegmayer *et al*., 2012; Milone *et al*., 2013). SOMs can represent complex high-dimensional input patterns into a simpler low-dimensional discrete map, with prototype vectors (neurons) that can be visualized in a two-dimensional lattice structure and which preserve the proximity relationships of the original samples (Kohonen *et al*., 2001). Interestingly, SOM has several advantages regarding other current methods (DiLeo *et al*., 2011; Chai *et al*., 2021). Indeed, SOMs looks at each data sample individually, it clusters genes with highly similar expression patterns along the features measured, and allows an individual, rather than a global, analysis of each gene/phenotype. In plant studies, SOMs have been widely used for the integration of transcriptome profiles, metabolites, the elucidation of gene-to-gene and metabolite-to-gene pathways (Allen *et al*., 2010; Hirai *et al*., 2005; Yokota Hirai *et al*., 2004; Yano *et al*., 2006), and for integration and discovery of coordinated variations in transcriptomics and metabolomics data (Stegmayer *et al*., 2009; Milone *et al*., 2010; Lopez *et al*., 2015).

In this study, we aim to identify genes as potential drivers for the establishment of Kranz anatomy. For that, expression data from Prochetto *et al*. (2023) was combined with anatomical information extracted from a detailed study of the leaf gradient in four non-model grass species from the Otachyriinae subtribe. We focused on three main components of leaf anatomy: (1) BS area, (2) patterns of venation and (3) connection between M and BS cells. Then, anatomical data and transcriptome information were combined using unsupervised ML methods to identify a set of transcription factors that could be responsible for the differentiation and evolution of C_4_ Kranz anatomy.

## Materials and methods

### Species selection

To carry out anatomical studies, four species of the Otachyriinae subtribe were selected: *Hymenachne amplexicaulis* (Rudge) Nees (C_3_), *Rugoloa pilosa* (Sw.) Zuloaga (PK), S*teinchisma hians* (Elliott) Nash (C_2_) and *Anthaenantia lanata* (Kunt) Benth (C_4_). Species selection was based mainly on phylogenetic proximity, photosynthetic pathway, and material availability. Seeds or rhizomes of the studied species were collected in the field. The specimens were deposited in the herbaria Instituto Botánica Darwinion (SI) and Arturo Ragonese (SF). The collection vouchers are listed below: *Hymenachne amplexicaulis* (Rudge) Nees Prochetto and Reinheimer 1 (SF), *Rugoloa pilosa* (Sw.) Zuloaga s / n (SI), *Steinchisma hians* (Elliott) Nash Marino s / n (SF), *Anthaenantia lanata* (Kunt) Benth Acosta (SI).

### Plant growth conditions and sampling

Individuals of four Otachyriinae species were grown, from seed or rhizome, in growth chambers at 27°C under long-day conditions (16 hours of light and 8 hours of darkness). For each species, the 5th young leaf of 10 individuals was collected. To study the leaf development gradient, each leaf was divided into two sections, taking the sink-source transition zone as reference. The sections were then divided into four segments of equal length and labeled S1 to S8 from the base to the tip of the leaf. Segments 1, 3, 5 and 7 were used for the analysis. To account for high variability between samples, replicates were paired throughout the analysis.

### Preparation and analysis of samples for light microscopy

Fresh fragments were placed in plugs with 5% low melting point agarose in 0.05 M PBS at 50°C and left until solidification at 4°C for at least 1 hour. Cross sections of 100 μm thickness were obtained with a vibrating blade microtome (Leica VT1000 S) and mounted on slides with 0.05 M PBS. Sections were visualized and photographed under fluorescence and white light microscopy (Nikon Eclipse E200). Image processing and anatomical measurements were performed using FIJI v2.9.0 (Schindelin *et al*., 2012).

### Statistical analysis

To determine the significance between the differences in the phenotypic characters, non-parametric methods were used contemplating paired samples (for intra-species comparisons) and unpaired (for comparisons between species) with Graphpad Prism (v6.0.1 for Windows, www.graphpad.com). For the paired samples, Friedman test and Dunn’s multiple comparisons test were used. For unpaired samples, Kruskal-Wallis test and Dunn’s multiple comparisons test were used. P-values lower than 0.05 were considered significant.

### Principal Component Analysis

Principal component analysis (PCA) was conducted in R, using stats package (v3.6.2, R Core Team 2021). Biplot graphs were generated using factoextra (v1.07, Kassambara and Mundt, 2022).

### Gene expression data

Transcriptomic data including gene annotation and transcripts abundance was retrieved from previous study (Prochetto *et al*., 2023). Gene Expression Matrices were built using the abundance_estimates_to_matrix.pl script from Trinity package v.2.8.5 (Haas *et al*., 2013), to generate a TPM expression matrix. Sequencing depth normalization was performed using the trimmed mean of M-values (TMM) method from the EdgerR package v. 3.38.1 (Robinson *et al*., 2009). The total number of genes used in this study was 9,739.

### Self-Organizing Maps

The goal of Self-Organizing Maps (SOMs) (Kohonen, 1982; Wehrens and Buydens, 2007) is to represent complex high-dimensional input patterns into a simpler low-dimensional discrete map, with prototype vectors (centroids of the neurons) that can be located in a two-dimensional lattice structure, while preserving the proximity relationships of the original data (Stegmayer *et al*., 2009; Milone *et al*., 2010). In this work we have developed two SOMs for different purposes. The first model was developed for grouping features (that is, genes together with phenotypic traits) into neurons according to their expression along leaf development and species with different photosynthetic subtypes. To this end, scaled expression values (log transformed, mean centered and variance-scaled) of a total of 9,757 features for all the three species under analysis were used. SOM training was made with aweSOM package (v 1.3., Boelaert *et al*., 2022), using an hexagonal grid of 289 neurons (17×17), rlen = 1000, alpha = c (0.1, 0.001) and Euclidean distance function. For each SOM, its corresponding quantization error, a parameter that assesses the accuracy of the SOM for representing the data, was calculated with the somQuality function from the aweSOM package (v1.3, Boelaert *et al*., 2022). The median value of the features grouped in neurons was used to characterize each neuron pattern, and thus, to analyze similarity between neurons.

The second model was a species-specific SOM built in order to focus on expression and phenotypic patterns of variation instead of their absolute expression magnitude; and to identify features that vary between species. Expression values were mean centered and variance-scaled separately for each species. Then C_3_ data (only) was used to train a 3 × 2 hexagonal SOM. This small size for the map was chosen in order to have a clear and easy way to analyze patterns of variation along leaf development in each neuron, as suggested in (Nakayama *et al*., 2018). Later, data from PK and C_4_ species was mapped to the species-specific SOM trained with C_3_ data. To visualize (as a directed network) the assignment of features from different species to separate neurons, igraph package v1.3.5 (Csardi *et al*., 2006) was used. Clustered and displaced feature sets among clusters were subjected to gene ontology (GO) analysis.

### Gene Ontology enrichment analysis

The Gene Ontology (GO) enrichment analysis was performed using the R package TopGO v2.42 (Alexa *et al*., 2022). GO terms for each transcript was obtained from Prochetto *et al*., (2023) annotation matrices. P-values were adjusted using the “elim’’ algorithm. A term was significant if its adjusted p-value < 0.01. To characterize neurons, GO BP terms were extracted and summarized with REVIGO tool (http://revigo.irb.hr, Supek *et al*., 2011).

## Results

### Leaf gradient anatomy characterization

We previously confirmed that the leaves of *H. amplexicaulis* and *R. pilosa* have the typical C_3_ anatomy whereas, *A. lanata* leaf presents C_4_ characteristics (Prochetto *et al*., 2023). Based on light microscope photographs and RNAseq data we identified segments 1 and 3 (S1, S3) as the sink zone of the leaf and segments 5 and 7 (S5, S7) as the source zone (Prochetto *et al*., 2023). In addition, differential expression analysis demonstrated the existence of a leaf developmental gradient in *H. amplexicaulis*, *R. pilosa* and *A. lanata* as was previously reported in several monocot species (Li *et al*., 2010; Pick *et al*., 2011; Wang *et al*., 2014; Studer *et al*., 2016; Prochetto *et al*., 2022).

To supplement available anatomical information on Otachyriinae species, we analyzed the anatomy and development of *S. hians* (C_2_) leaf and compared them with additional anatomical observations in *H. amplexicaulis* (C_3_), *R. pilosa* (PK) and *A. lanata* (C_4_) (Fig. 1; Supplementary Fig. S1). Definitions and units of phenotypic traits are shown in Supplementary Table S1. Traits that did not show significant differences among the leaf segments, despite having an observable developmental pattern throughout the leaf, are presented in Supplementary Fig. S2. Traits that showed significant differences (P-values lower than 0.05) among leaf segments and species are summarized in Fig. 2A.

**Figure 1:**
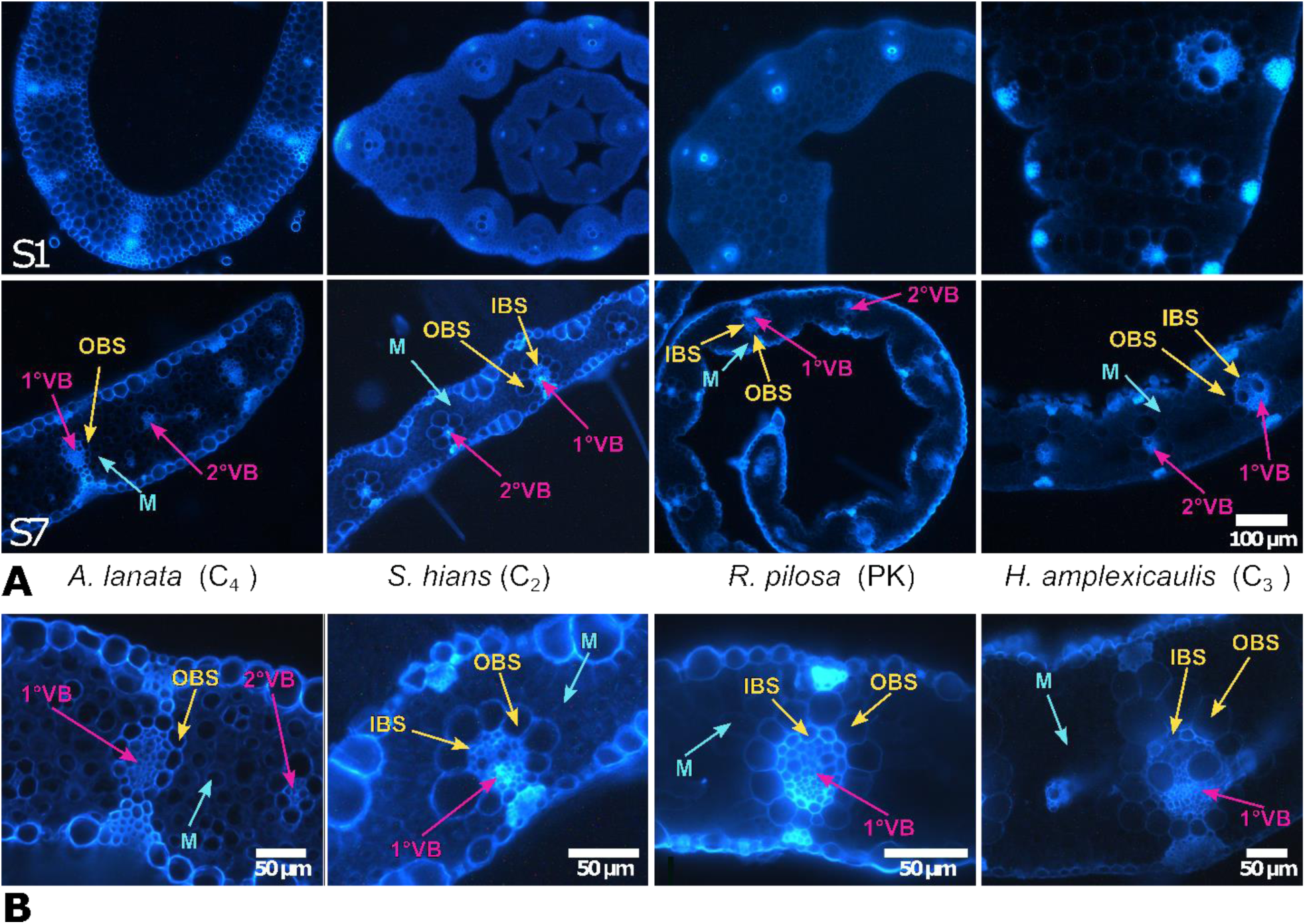
Developmental gradient in the 5th leaf of four Otachyriinae subtribe species. (A) S1 and S7 cross section cuts from C_4_ *A. lanata*, C_2_ *S. hians*, PK *R. pilosa* and C_3_ *H. amplexicaulis*. (B) Magnification showing 1° vascular bundles from S7. Abbreviations: Primary vascular bundle (1°VB), Secondary vascular bundle (2°VB), Inner Bundle sheath cell (IBS), Outer bundle sheath cell (OBS) and Mesophyll cell (M).

**Figure 2:**
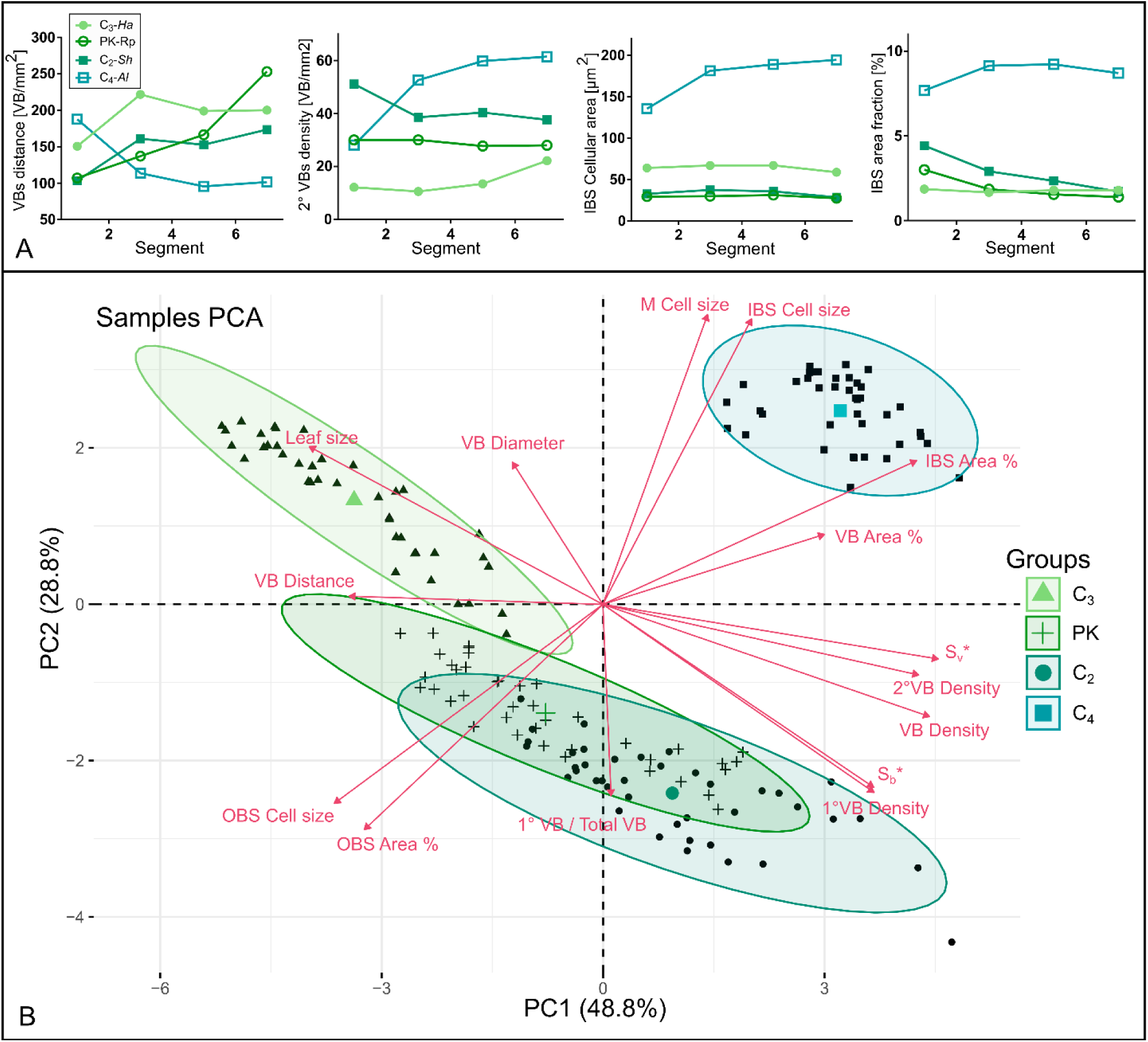
Summary of leaf development traits. (A) Phenotypic traits along the leaf gradient with divergent patterns among species. (B) Principal component analysis of the samples using phenotypic traits (PCA biplot). Abbreviations: Mesophyll cells (M), Vascular Bundle (VB), Inner Bundle Sheath cells (IBS), Outer bundle sheath cells (OBS).

### Leaf development traits associated with vascular bundles

The vascular bundles (VB), placed next to the Inner and Outer Bundle Sheath cells (IBS, OBS), play a fundamental role in the structure of the leaf anatomy. In this section, we study how vascular bundle traits vary throughout development and how these patterns differ in species with different photosynthetic subtypes. For that, we analyzed: (a) distance between central VB and its closest VB, (b) the number of VB per cross section area, (c) the number of 1° VB per cross section area, (d) the number of 2° VB per cross section area, (e) proportion of 1°VB, (f) mean VB diameter, and (g) proportion of cross section area occupied by VB.

The distance between VB tends to increase throughout the leaf development of the C_3_, PK and C_2_ species, while it decreases in the C_4_ species (Fig. 2A). This trend generates a decrease in VB density as leaves mature in PK and C_2_ and an increase of VB density in C_4_ (Fig. 2A). This trend is not clear in C_3_ where the VB distance increases in S3 and then remains without major changes (Fig. 2A). In terms of VB distance and VB density we observed that the most remarkable changes happen between S1-S3 in all the studied species.

When the cross-sectional area of vascular bundles (VB Area %; Supplementary Fig. S2) was analyzed, we observed a descending pattern from S1 to S7 for all species. This trend is, significantly, pronounced in C_2_ and C_4_ species. This phenomenon is explained both by the reduction in the diameter of the primary VBs and the appearance of new smaller secondary VBs (Supplementary Fig. S2).

### Leaf development traits associated with bundle sheath and the mesophyll cells

Among the most relevant characteristics of the Kranz anatomy are the larger area of bundle sheath cells (BS) and their special arrangement around VB (Esau, 1953). To quantify characteristics of Kranz anatomy, we selected eight traits to document the area of the IBS, OBS and mesophyll (M) cells and the fraction of tissue they occupy within the leaf cross section. These traits included: (a) mean IBS area in the cross section, (b) proportion of cross section area occupied by IBS, (c) mean OBS area in the cross section, (d) proportion of cross section area occupied by OBS, (e) mean M area in the cross section, (f) cell interface between BS and M per cross section area (S_b_*), (g) cell interface between VB and BS per cross section area (S_v_*), and (h) total cross section area of the leaf.

In the C_3_ species, both the IBS area and the fraction it occupies in the tissue are constant along leaf developmental gradient (Fig. 2A). In PK, no differences were observed in the IBS area; however, we observed a decrease in the fraction of the IBS area as leaves mature (Fig. 2A). A similar trend is observed in C_2_ (Fig. 2A); nevertheless, this pattern is accompanied by an increase in the IBS area in segments S3 and S5 (Fig. 2A; Supplementary Fig. S2). In the C_4_ species, an initial and significant increase in the area of the IBS is observed; however, it does not result in significant changes in the fraction occupied by the IBS throughout development (Fig. 2A; Supplementary Fig. S2).

In all C_3_, PK and C_2_, at S3 and S5, the OBS increase in cell area and in the fraction that they occupy (Supplementary Fig. S2). While cell area of IBS cells increases from base to tip (e.g., C_2_ and C_4_), the average area of the mesophyll cells is constant along the leaf (Fig. 2A; Supplementary Fig. S2).

A Principal Component Analysis (PCA) of samples, based on phenotypic traits, showed a clear distinction between C_4_ and the rest of the species, with some overlap between PK and C_2_ species (Fig. 2B). Principal Component 1 explains 48.8% of the variation and Principal Component 2 explains 28.8 1% of variation. The phenotypic traits contributions to the distribution of the samples are shown as vectors in the biplot. We observed that M cell area, IBS area and to a lesser extent VB area %, are the main traits that differ between C_4_ species and the rest (Fig. 2B). On the other hand, traits related to VB density and contact areas (S_b_*; S_v_*) are positively correlated with the shift from C_3_ to C_2_.

### Comparison of phenotypic traits between species

Traits associated with VB, BS, and interfaces between cell types were compared in mature segments of the three sampled species (Fig. 3). The distance between VB of C_4_ was the shortest among species resulting in a higher density of VB. Overall, a crescent VB density is observed from the C_3_ species through PK species and, successively, towards C_2_ and C_4_ (Fig. 3A). We observed that the proportion of primary VB over the total of VB was significantly smaller in C_4_ (Fig. 3A). Leaf fraction occupied by IBS was significantly higher in C_4_ (Fig. 3B). Similarly, the leaf fraction occupied by OBS was significantly higher in C_2_ and lower in PK (Fig. 3B).

**Figure 3:**
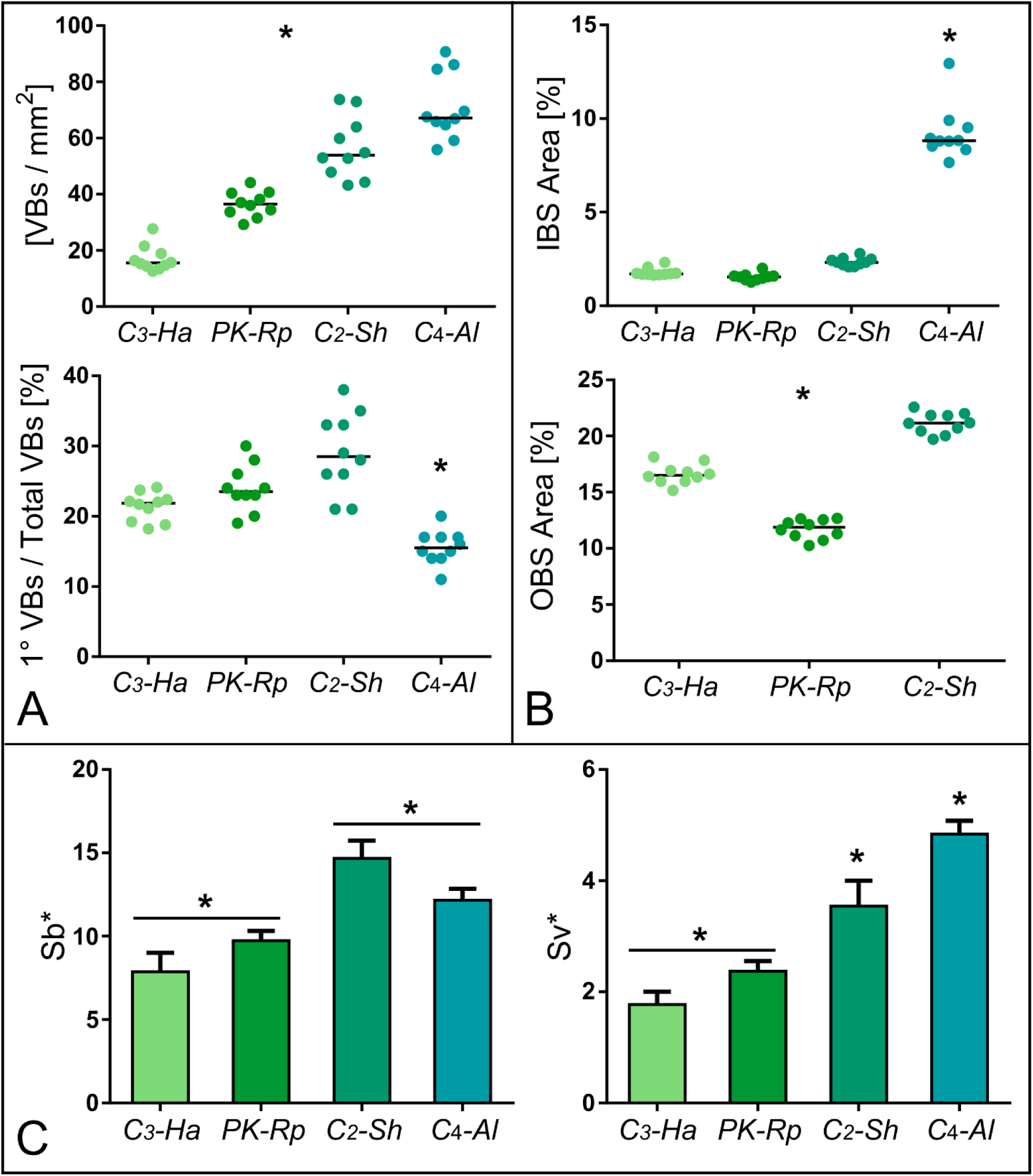
Phenotypic traits comparison between species. (A) Phenotypic traits associated with vascular bundles. (B) Phenotypic traits associated with bundle sheath cells. (C) Parameters associated with interfaces between cell types. Asterisks point out statistical differences between groups (Kruskal-Wallis test and Dunn’s multiple comparisons test, p value < 0.05). Abbreviations: Mesophyll cells (M), Vascular Bundle (VB), Inner Bundle Sheath cells (IBS), Outer bundle sheath cells (OBS).

The need to maintain a significant transport of metabolites between BS and M in the C_4_ and C_2_ species suggests the need for an anatomical adaptation that facilitates transport. In addition to increasing the density and size of the plasmodesmata, this can be achieved by increasing the contact surface between both cell types. The surface parameter S_b_, defined by Pengelly *et al*. (2010), is calculated on leaf cross section and by measuring the perimeter of the BS within the space between two VB and dividing it by the distance between VB. However, this parameter does not consider the nonlinear organization of the VB in *A. lanata*. So, to include the tridimensional organization of VB in Pengellýs equation we here proposed a variation of this parameter (S_b_*). The S_b_* parameter is calculated as the perimeter of all the BS cells in contact with the mesophyll divided by the cross-sectional area of the leaf. The results showed that C_2_ and C_4_ S_b_* are significantly larger than the C_3_ species (Fig. 3C).

We also estimated the contact surface between BS and VB. For that, we calculated the perimeter of all the BS cells in contact with the VB divided by the cross-sectional area of the leaf (S_v_*). The results showed that C_2_ and C_4_ have S_v_* significantly larger than the C_3_ species (Fig. 3C).

### SOM optimization for the integration of phenotypic traits with gene expression data

The expression data from Orthogroups (OG) from three species (*A. lanata*, *R. pilosa* and *H. amplexicaulis*) and four segments of leaf gradient (S1, S3, S5 and S7) (Prochetto *et al*., 2023) were merged with phenotypic trait data to build a SOM. This first SOM allows us to identify subsets of genes with similar expression profiles and link them with the behavior of phenotypic traits along the leaf gradient from different species.

Before building this SOM, the optimum grid size was studied in the following way. In order to make the analysis of the results of the individual neurons affordable, a median neuron size from 32 to 37 features per neuron was set (Supplementary Table S2). The total number of features (genes and phenotypic traits) measured for all three species was 13,953. By using different cut-off expression values as a threshold for the genes, several subsets were obtained (ranging from 13,953 to 2,804 according to the median number of features per neuron desired) and used to build SOMs of different sizes (from 400 to 81 neurons in total). The minimum quantization error was achieved for the SOM with 9,757 features and 289 neurons (17×17 map) with a median number of features per neuron of 32 (Supplementary Table S2) (Fig. 4). The selected SOM includes 9,739 genes and 15 phenotypic traits and three species-specific features (Supplementary dataset 1). To explore grouped features neuron by neuron an interactive version of the SOM is shown in Supplementary Fig. S3. Overall, transcription factors and genes involved in cell wall development are widely distributed along the map. Meanwhile, some classes like photosynthesis and phenotypic traits are concentrated in more specific areas. When the median values of the grouped features were analyzed, we observed, as expected, that close neurons present similar patterns along the samples (Supplementary Fig. S4).

**Figure 4:**
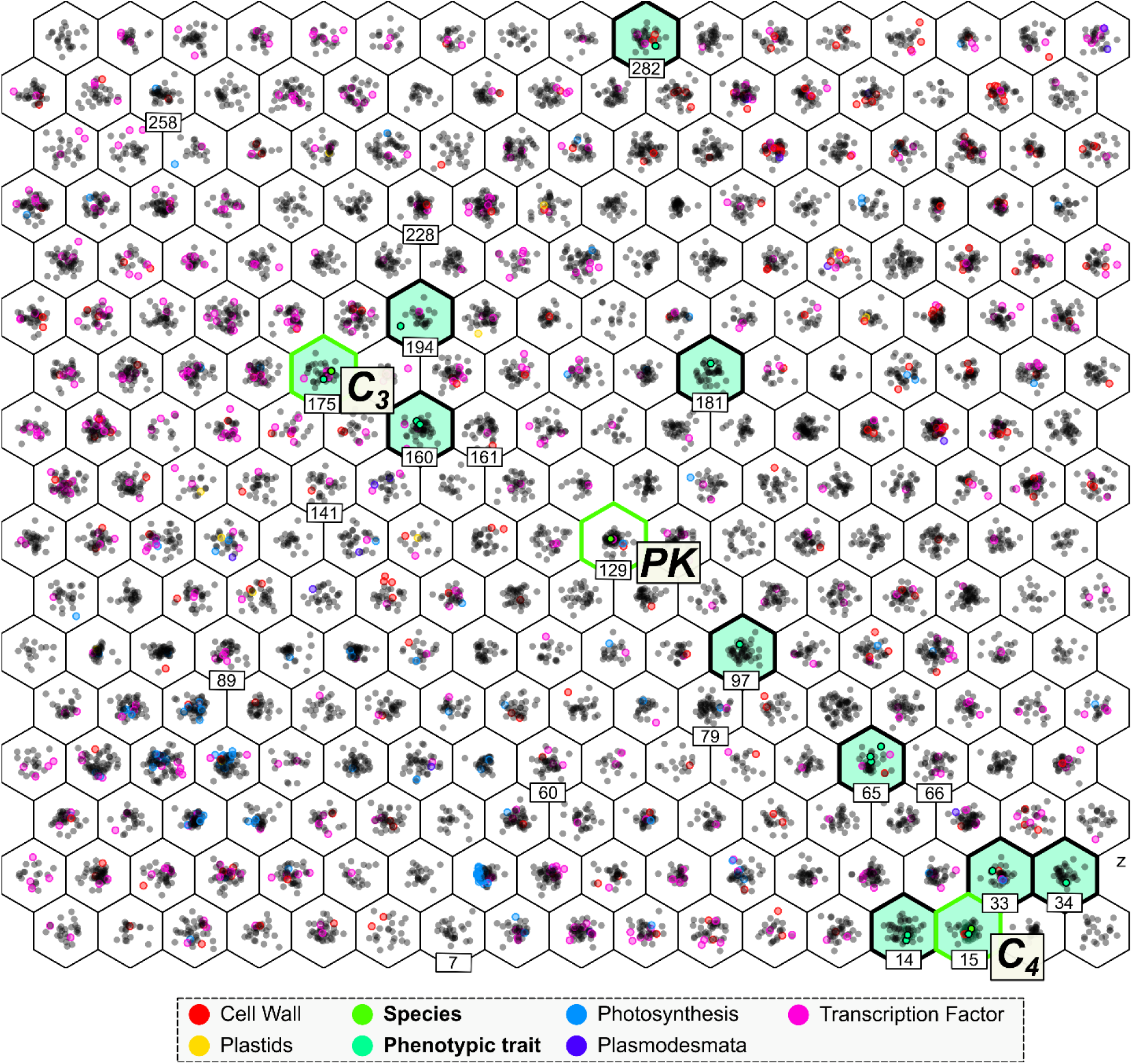
Self-Organizing Map from Supplementary dataset 1 (with a total of 9757 features coming from leaf phenotypic information (18 features) and gene expression data (9739 genes) for C_4_ *A. lanata*, PK *R. pilosa* and C_3_ *H. amplexicaulis*. Each cell represents a neuron and each point a feature. Neurons containing phenotypic traits are highlighted in green. Different types of features are highlighted in colors.

### GO terms in selected neuron pairs

Considering that SOM only considers positive correlation among features, we looked for pairs of neurons with opposite patterns in order to find neurons that better represent both types (positive and negative) of potential relationships between genes and phenotypic traits. Pairs of neurons having more than one phenotypic trait and opposite patterns associated with GO terms are shown in Fig. 5. A detailed membership list of each neuron pair is shown in Supplementary dataset 2.

**Figure 5:**
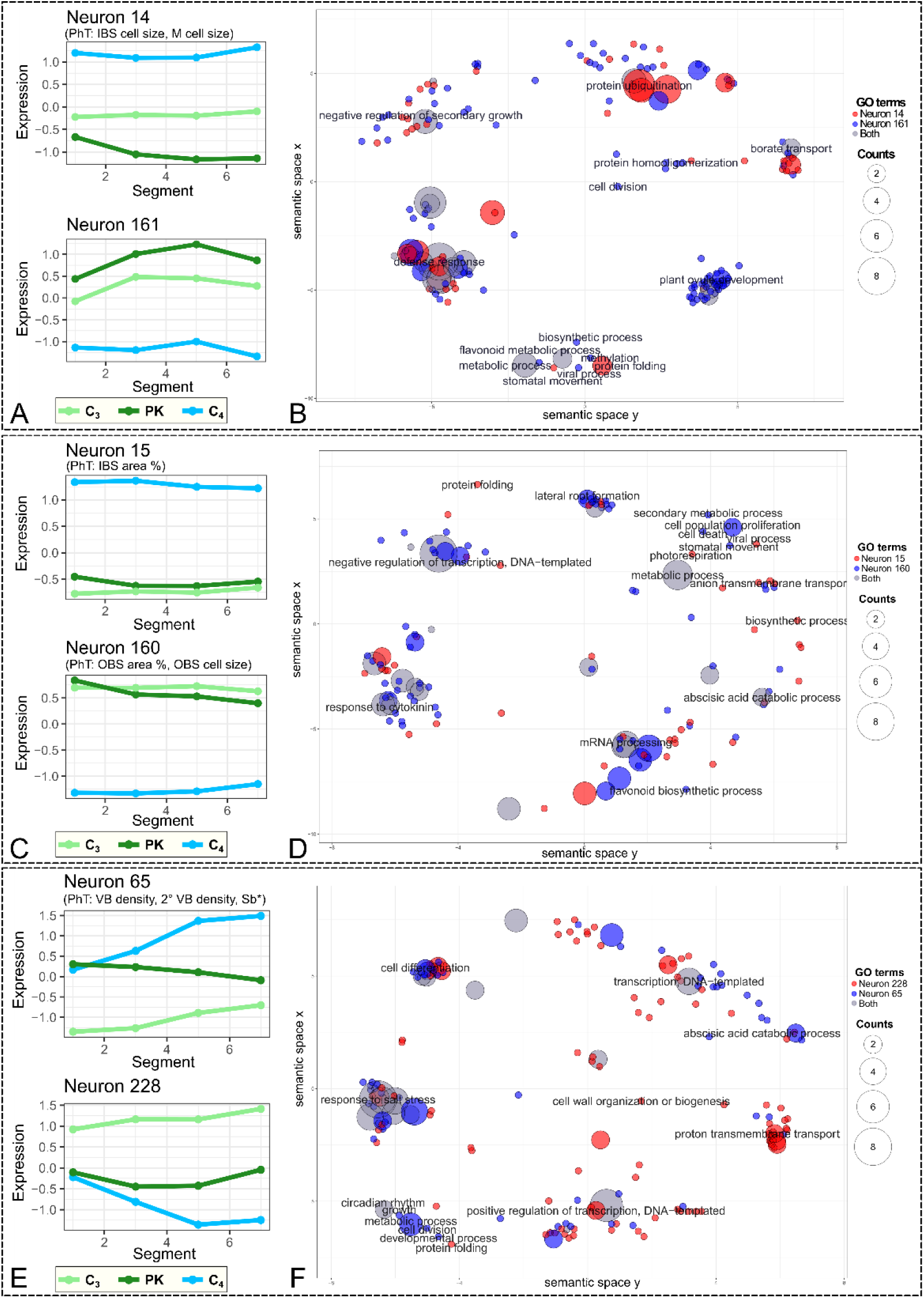
Selected pairs of opposing neurons containing phenotypic features: N14 and N161. (A, B), N15 and N160 (C, D) and N65 and N228 (E, F). Neuron median expression values along samples (A,C,E). Semantic similarity clustering of gene ontology terms (BP) present in neuron pairs made by Revigo (B, D, F). Circle colors indicate if a term is present in one neuron (red and blue) or in both neurons in the pair (gray). Circle size indicates the times in logarithmic scale that a term is present in the reference database. Abbreviation: PhT, phenotypic trait.

Phenotypic traits IBS cell area and M cell area were in neuron N14 from the N14-N161 pair (Fig. 5A). N14 contains features with high expression in C_4_ and low expression in PK, with C_3_ in between. A wide range of processes represented as GO terms clusters were present in the pair (Fig. 5B), including defense response, protein ubiquitination, plant ovule development and regulation of secondary growth as the most numerous clusters. Although most of the terms were present only in one of the neuron pairs, most of the clusters presented terms from both neurons (Supplementary Fig. S5). This pair also contains one transcription factor (OG0011232, inside N161) belonging to the NF-YA family.

IBS and OBS area fraction (%) and OBS cell area were in the N15-N160 pair (Fig. 5C). N15 features show high values in C_3_ and PK species and low values in C_4_, meanwhile neuron 160 shows the opposite pattern. GO terms present in the pair were clustered in 16 clusters, but 10 of them were singletons (Fig. 5D; Supplementary Fig. S6). The remaining six concentrate most GO terms and were related to response to cytokinin, mRNA processing, negative regulation of transcription, lateral root formation, abscisic acid catabolic process and anion transmembrane transport. Three transcription factors from three different TF families were present in N160: OG0003129, a member of C3H zinc finger family; OG0004661, a member of HD-ZIP family; and OG0009193, belonging to GRAS family.

VB density, 2° VB density and S_b_* phenotypic traits were assigned to N65, belonging to the N65-N228 pair (Fig. 5E). Neuron 65 features show increasing values along the leaf gradient for C_4_ species, and from C_3_ to C_4_ species meanwhile neuron 228 shows the inverse pattern. GO terms present in the pair were clustered in 13 clusters (Fig. 5F; Supplementary Fig. S7), with seven of them being single term clusters. The remaining six clusters were involved in response to salt stress, transcription and regulation of transcription, cell differentiation, proton transmembrane transport and abscisic acid catabolic process. The pair also included five transcription factors, two inside N65 (OG0005274, OG0010188) and three inside N228 (OG0001295, OG0004381, OG0006982).

### Species-specific SOM for studying conservation of expression and phenotypic patterns of variation across evolution

To study the changes in features (genes and phenotypic traits) expression patterns associated with photosynthetic evolution, a species-specific SOM was built with the C_3_ data alone. The grid size was chosen to be small (3×2) so that neurons represent unique and non-redundant expression patterns which can capture the behavior of features along the leaf gradient of our dataset (Supplementary Fig. S8). Then we used the C_3_ species-specific trained SOM to map PK and C_4_ data to identify two different scenarios: (a) features that are preserved in the same neurons among species and (b) features that are allocated in different neurons among species. Indeed, a feature with a very similar expression pattern across different species will be always mapped to the same neuron in the trained map, meaning that the feature is preserved across all species in this study. In contrast, a feature mapped to a different neuron means that the feature expression pattern is different between species, and therefore the feature will be displaced (from a particular neuron in species A to another neuron in species B). The analysis performed here assumes that feature preservation status gives information on the preservation of gene regulatory networks between species. To summarize the results, we have built a network graph to show features allocation to neurons and displacements from C_3_ to PK (Fig. 6A) and from PK to C_4_ (Fig. 6B). Furthermore, the quality of the displacement was assessed by inferring the relationships between the neuron expression patterns. Based on the results we identified four groups of expression pattern displacements: early, when it looks shifted to more immature segments in the leaf gradient; delay, when the displacement looks shifted to more mature segments in the leaf gradient; flip, when it presents the opposite pattern; and others, that groups the rest of expression pattern displacements with no specific meaning.

**Figure 6:**
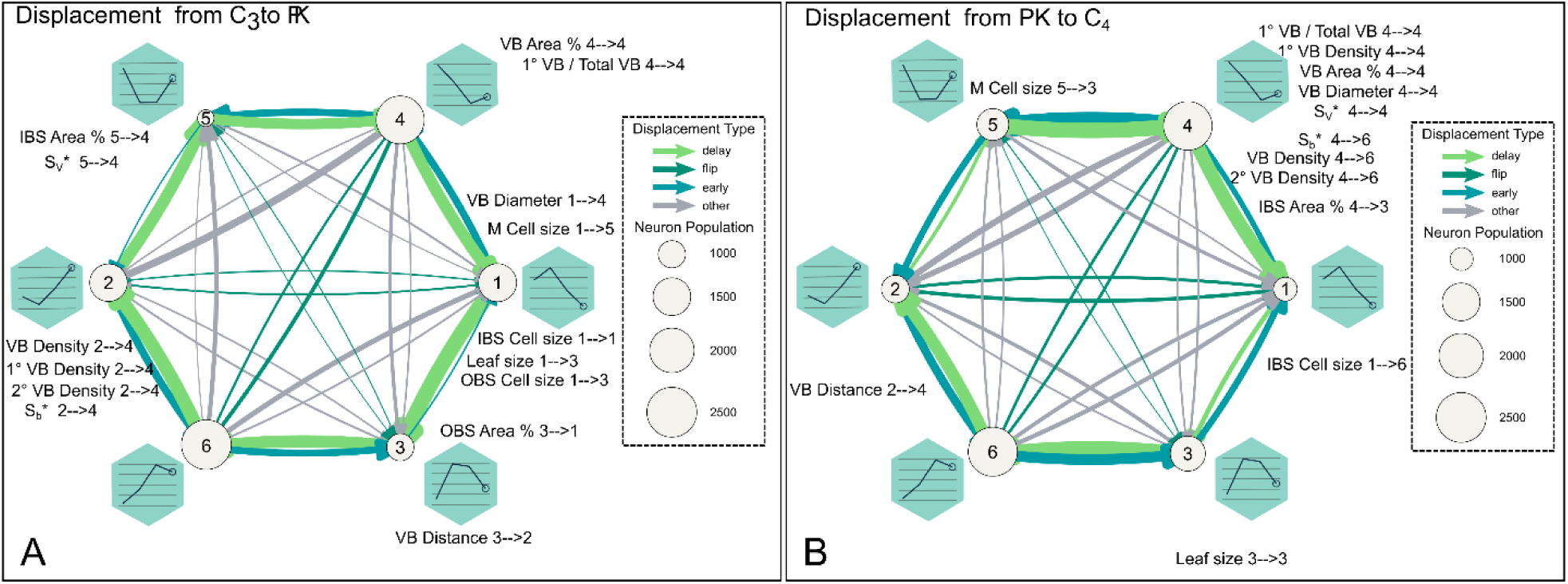
Displacement of features in different clusters in the SOM clustering scheme. Network representations of feature assignment into different SOM clusters using the displaced features. Arrows represent displacement from C_3_ to PK (A) and from PK to C_4_ (B). Arrows and circle sizes are proportional to the number of displaced features and neuron population, respectively. Line plots indicate representative expression patterns (neuron median expression) in each cluster throughout the leaf gradient. Arrow colors show distinct displacement types.

A total of 5,469 features were displaced from C_3_ to PK (Fig. 6A, Supplementary Fig. S9). Almost 70% of displaced features were displaced to a close neuron (3,786 features), either to an earlier (1,145 features, 20.9%) or delayed (2,641, 47.7%) expression pattern. Only 438 features (8.0%) presented a flip displacement, while other kinds of displacements altogether accounted for 1,245 features (22.7%). Only three of the phenotypic traits were preserved between C_3_ and PK (IBS cell area, VB area % and 1°VB/Total VB). The early group of displaced features contains only one phenotypic trait (VB diameter) and is enriched in carboxylic acid catabolic process, response to stress and leaf development, among others (Supplementary dataset 3). Two phenotypic traits belong to the delay group of displaced features (leaf area and OBS cell area). A GO enrichment analysis showed that this group is significantly enriched in RNA metabolic processes (Supplementary dataset 3).

A total of 5,544 features were displaced from PK to C_4_, with most of them displaced to a close neuron (67.7%) as in the C_3_ to PK transition (Figure 6B, Supplementary Fig. S9). However, these account for a higher number of early displacements (1,687, 30.4%) and a lower number of delay displacements (2,068, 37.3%) than in C_3_ to PK transition. Also, while the delay displacements show a homogenous distribution across neurons between C_3_ and PK, in the PK to C_4_ transition, some displacements, as from neuron 1 to 3 or neuron 2 to 5 are rare. A higher number of phenotypic traits (leaf area, VB area %, 1°VB/Total VB, VB diameter, 1° VB density and S_v_*) were preserved in this transition than in the C_3_-PK transition. A GO enrichment analysis showed that the early group of displaced features is enriched in mRNA processing and seed germination, among others (Supplementary dataset 3). The delay features from PK to C_4_ are significantly enriched in developmental processes involved in reproduction, proteolysis, cell morphogenesis and developmental cell growth among others (Supplementary dataset 3).

### Genes displaced together with specific phenotypic traits

Addressing the displacement of phenotypic traits between species is another way of identifying potential regulators involved in photosynthetic pathway evolution. Overall, there is not a clear trend in the type of phenotypic traits displacements (Supplementary Fig. S10). The area of the BS is an interesting trait postulated as one of the most important in Kranz anatomy, because it is where CO_2_ carboxylation takes place in C_4_ species (Christin *et al*., 2013, Lauterbach *et al*., 2019). Our results show that the IBS cell area is preserved between C_3_ and PK species (Fig. 6A, both belonging to neuron 1) but it is displaced to neuron 6 in C_4_ species (Fig. 6B). The IBS expression changes from one descending pattern from base to tip, to an ascendant pattern in the C_4_. We found that together with this trait there are also 41 genes displaced. A GO enrichment analysis showed that this group of genes is significantly enriched in suberin biosynthetic process, pyruvate transport and cell wall modification, among others (Supplementary dataset 4). Indeed, two genes in this group are known to be part of suberin biosynthesis; OG0000296, an ortholog to AtFAR1/4 (Fatty Acid Reductase) and OG0005527, an ortholog to AtABCG16, a class of ABCG half-transporters. In addition, there are two transcription factors among the features displaced. One is a member of the Class 1 TCP transcription factor family (OG0009002, AtTCP7). The second one (OG0012289, AtBFP4) is a member of the GeBP family.

## Discussion

A previous study on 157 closely-related grass species demonstrated that only a few phenotypic traits had significant differences between C_3_ and C_4_ species (fraction of the leaf occupied by OBS and IBS, M cell number between VB and IBS cell area) (Christin *et al*., 2013). Overall, more differences were observed between C_3_ species of the BEP (sometimes BOP clade) and PACMAD clade than between C_3_ and C_4_ species of the PACMAD clade, where the values showed a tendency to overlap (Christin *et al*., 2013). In dicots, Lauterbach *et al*. (2019) found a similar trend in the Tribuloideae tribe (Zygophyllaceae) (Lauterbach *et al*., 2019). In that work, it was observed that the C_4_ species of the tribe have a greater fraction of the area occupied by the BS. Notably, they also observed that the C_3_ species in evolutionary proximity to the C_4_ species have an area occupied by BS with similar dimensions to the C_4_ species. Both works suggest that many of the anatomical features, such as the distance between VB, cell areas and the area occupied by BS, are ancestral features on the evolutionary path towards the establishment of a Kranz anatomy and C_4_ photosynthesis.

Furthermore, it was found that the VB density was only significantly higher for C_4_ species in relation to C_3_ species (Lauterbach *et al*., 2019). In the same sense, Lundgren *et al*. (2019) established that, in *Alloteropsis semialata* (Poaceae), the only trait shared by all C_4_ populations was a higher VB density (Lundgren *et al*., 2019). These results demonstrate that the increase in VB density would not be a necessary anatomical preconditioner to drive the change toward a C_4_ anatomy, as previously postulated (Sage *et al*., 2014). These works highlight the importance of the selection of species and their evolutionary context in trying to establish important phenotypic markers for the evolution of C_4_ photosynthesis.

### There are structural differences in the leaf anatomy from the beginning of the development gradient

The study of the leaf anatomy presented in this work demonstrated the existence of a leaf development gradient. Overall, we found conserved and non-conserved patterns of leaf anatomical development among the studied species. In particular, the most significant phenotypic changes occurred between S1 and S3, except for the fraction of tissue occupied by IBS and the area of the M cells. The PK and C_2_ leaves were most similar in terms of leaf anatomy; however, as expected, the C_2_ leaves presented a higher degree of Kranz characteristics such as density of the VB, fraction occupied by the OBS and the contact surfaces between the different cell types.

### Communication between BS and M as a possible indicator of the Kranz degree of species

It has been suggested that a higher fraction occupied by BS in C_4_ species compared to C_3_ species is one of the distinguishing features of C_4_ anatomy (Sage, 2017; Christin *et al*., 2013; Lauterbach *et al*., 2019). Some of the C_4_ species have two Kranz compartments, OBS and IBS; however, most of the C_4_ species have only one. In this work, anatomical comparisons were made using a C_4_ species of the NADP-ME sub-type that has IBS. To date, there is not enough evidence to identify the moment in which natural selection opted for IBS or OBS as the Kranz compartment and the moment and reasons why C_4_ NADP-ME species lack OBS. Concerning this, it is thought that C_4_ species that use IBS as a Kranz compartment come from a C_4_ ancestor with OBS (Christin *et al*., 2013). In this work, we determined that in the C_4_ species, the IBS are up to 5.8 times larger than in other species, but OBS in non-C_4_ species are 2 to 8 times larger than IBS in C_4_. However, the fraction occupied by the IBS (IBS Area %) of the C_4_ species is smaller than the OBS fractions (OBS Area %) of the C_3_ species. These results agree with previous works (Christin *et al*., 2013; Lauterbach *et al*., 2019).

For many years, it has been thought that vein density may be an important precondition to acquiring the C_4_ pathway at the macroevolution scale. However, recent works suggested that VB density may not be a necessary precondition of a C_3_ ancestor to promote the evolution of C_4_ photosynthesis, but rather a trait that evolved at a later stage (Lundgren *et al*., 2019; Lauterbach *et al*., 2019). In line with this, in our work, we found that traits associated with VBs are early conditioned by anatomical differences between species. Here we found that VB density is remarkably influenced by VB spatial configuration, 1°/2° ratio, and area.

The importance of the area of the BS as a Kranz trait is given by the assumption that the BS must be large enough to house many chloroplasts that perform the CBB cycle and thus maintain a good photosynthetic efficiency (Sage and Sage., 2004). However, recently, the importance of the transport of C_4_ acids through plasmodesmata between M and BS has been highlighted as a constraint for this process (Danila *et al*., 2016; Danila *et al*., 2018). Indeed, it has been shown that the C_4_ species have a higher density of plasmodesmata at the interfaces between BS and M, as well as a greater contact surface between both cell types, while the area of the BS is not significantly different between C_3_ and C_4_ species (Danila *et al*., 2016). In this scenario, the optimization of the anatomy in the PK condition could have had great significance for the evolution of C_2_ and C_4_, since a greater activity of GDC in the VB would allow a loss of activity of this enzyme in the M to be non-lethal (Sage *et al*., 2014). Later in evolution and once the GDC was allocated in the VB, the selection pressure could have changed: instead of having inhibitory effects, photorespiration would become a resource to supply CO_2_ to the VB chloroplasts. This would imply a selection pressure to generate an increase in the contact surface, an increase in plasmodesmata, and other mechanisms that make glycine exchange between M and VB more efficient (Sage *et al*., 2012). In line with this, overall, we observed an ascending pattern in the magnitude of the contact surface parameters from C_3_ to C_4_ species. Given the lack of information on plasmodesmata density in intermediate species here presented, the place of this event in the evolution of C_4_ photosynthesis in Otachyriinae is unknown. Remarkably, *S. hians* and *A. lanata* manage to increase the contact surface partly through different strategies. Although *S. hians* presents OBS with an area like those of *R. pilosa* and much smaller than that of *H. amplexicaulis*, its high contact surface is explained by an increase in the density of VB and a decrease in the number of M between the VB. Comparing *S. hians* with *A. lanata* we see that the latter has IBS that are less than half the area of the OBS of the former. However, *A. lanata* has a higher density of VB, in particular secondary VB, achieving a similar contact surface with smaller VB. A similar trend was observed in the genus Flaveria and grass species that use IBS as a Kranz compartment (Christin *et al*., 2013; McKown and Dengler, 2009; McKown and Dengler, 2010). The determination of the densities and areas of the plasmodesmata in these species would help to corroborate the dynamic of the interface between M and BS cells in Otachyriinae.

### Crossover data from transcriptomics and leaf anatomy shows potential candidate genes for the establishment of Kranz phenotypic traits

Deciphering gene network contributions to switches in phenotypic states, such as observed during leaf development and the evolution of photosynthesis, will entail identifying individual signaling gene factors as candidates for the establishment of the Kranz anatomy.

Using SOM we were able to identify transcription factors that correlated with particular phenotypic traits along leaf development and may be responsible for the evolution of leaf anatomy in Otachyriinae. A list of these potential gene drivers is shown in Table 1.

**Table 1:**
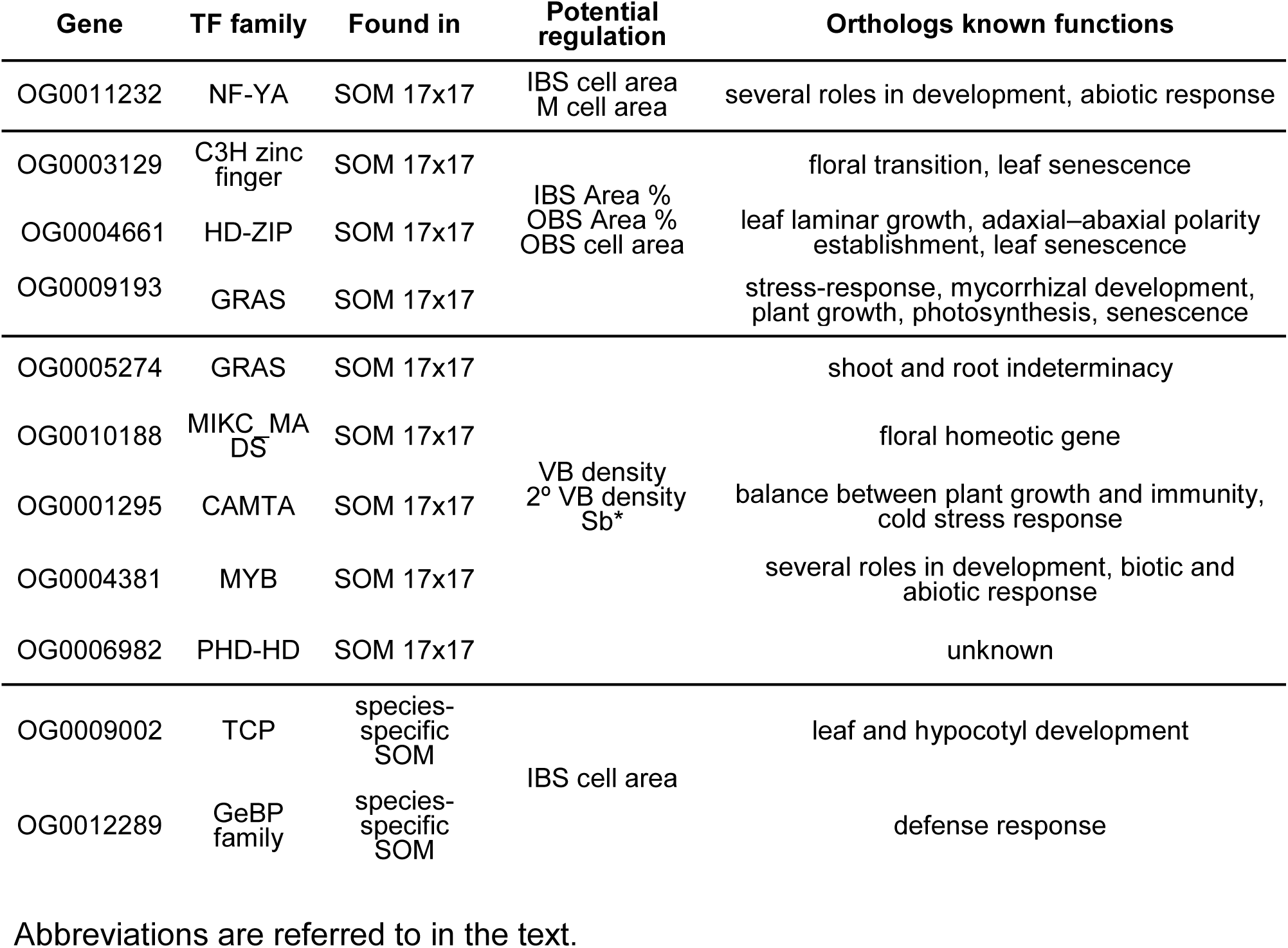
Transcription factors (TF) potentially involved in leaf phenotypic traits differentiation.

### Transcription factor correlated with the area of M and IBS cells in Otachyriinae

One transcription factor belonging to the NF-YA family was highly correlated with the area of the BS and M cells. NF-Y transcription factors are encoded by a single gene in animals and yeast. However, in plants, each NF-Y subunit type is encoded by 10 or more genes due to gene duplication events (Edwards *et al*., 1998; Siefers *et al*., 2008; Yang *et al*., 2016). Members of this family have been reported to play several roles during the life cycle of Arabidopsis and other dicot species (Populus and tomato) such as gametophyte development, embryogenesis, seed development, and post-germinative growth, abscisic acid (ABA) signal transduction, flowering cycle regulation, primary root elongation, blue light response and abiotic stresses (Mu *et al*., 2013; Yu *et al*., 2021; Hwang *et al*., 2019; Ballif *et al*., 2011; Warpeha *et al*., 2007; Li *et al*., 2021). In addition, it has been shown that the transcription factors CONSTANS and Nuclear transcription Factor Ys (NF-Ys) form a complex that governs light-dependent photoprotective responses in green alga *Chlamydomonas reinhardtii* promoting a photoprotective response in algal photosynthesis and flowering in plants (Tokutsu *et al*., 2019). In maize (*Zea mays*) and rice (*Oryza sativa*) members of the NF-YA regulates the expression of multiple genes related to stress resistance and growth, thereby improving the biotic and abiotic resistance of plants (Alam *et al*., 2015; Lee *et al*., 2015; Li *et al*., 2017; Wang *et al*., 2018; Lv *et al*., 2022; Tan *et al*., 2022). Additionally, ZmNFYA03 promotes early flowering by binding to the FT-like12 promoter in maize (Su *et al*., 2018).

### Transcription factors correlated with the area of IBS and OBS cells and area fraction of the leaf in Otachyriinae

We identified three transcription factors that correlate with the IBS and OBS area fraction and the OBS cell area. Such transcription factors belong to three different families: (1) C3H zinc finger family, (2) HD-ZIP family, and (3) GRAS family. The *A. thaliana* orthologs of C3H zinc finger family play a positive role in floral transition and leaf senescence (Yan *et al*., 2017). Aba insensibly to growth 1 (ABIG1), *A. thaliana* ortholog of HD-ZIP family plays a diverse role in leaf development such as leaf laminar growth, adaxial–abaxial polarity establishment, and leaf senescence (Preciado *et al*., 2021; Kollmer *et al*., 2015). The GRAS family gene member belongs to the LISC subfamily. The LISCL subfamily is involved in several processes such as stress response, mycorrhizal development, plant growth, photosynthesis, and senescence (Fode *et al*., 2008; Chen *et al*., 2015; Xu *et al*., 2015; Xue *et al*., 2015; Cenci and Rouard, 2017). In terms of photosynthesis, transcript analysis showed that *Haloxylon ammodendron* ortholog of AT2G29060 is a putative negative regulator in C_4_ assimilating shoots compared with the C_3_ cotyledons (Li *et al*., 2015). Expression atlas also showed moderate expression during root and leaf development for maize and rice OG0009193 orthologs (Dadvison *et al*., 2012; Stelpflug *et al*., 2016; Hoopes *et al*., 2018).

### Transcription factors correlated with the VB density and interface between M and BS cell cells in Otachyriinae

We identified five transcription factors that correlate with the VB density and interface between M and BS cells. Such transcription factors belong to five different families: (1) GRAS family, (2) MADS-box family, (3) CAMTA family, (4) MYB family, and (5) PHD-finger homeodomain family. The GRAS member belongs to the HAM subfamily that is implicated in shoot apical meristem maintenance and axillary meristem formation in *A. thaliana* (Schulze *et al*., 2010). The MADS-box family is well known to play key roles in flower development (Bowman *et al*., 1989; Coen & Meyerowitz, 1991; Weigel & Meyerowitz, 1994). Members of the CAMTA family are involved in drought stress tolerance, response to cold, and defense response in Arabidopsis (Zeng *et al*., 2022; Yuan *et al*., 2018). It has been documented that members of the MYB family play a role in the abiotic stress response and xylem development among others (Xiao *et al*., 2021; Wang *et al*., 2021). The PHD-finger homeodomain is a small family of transcription factors, whose functions are not yet very explored.

### Transcription factors correlated with the evolution from a C_3_ leaf anatomy to the C_4_ Kranz anatomy

Performing the species-specific SOM including all the phenotypic traits with transcriptomic data at once resulted in displacements without clear trends. Given these results, we found to better assess phenotypic traits individually. In this manner, we were able to look for the genes that are displaced together with a particular phenotypic trait. More specifically, we were interested in transcription regulators.

Overall, the number of features displaced from C_3_ to PK and PK to C_4_ was similar; however, the nature of such displaced features differs between C_3_/PK vs PK/C_4_. Features displaced from C_3_ to PK denoted a change in the moment of expression along development. Indeed, most of the displaced features showed a delay in their expression pattern. Interestingly, leaf area, S_v_*, IBS Area % and OBS cell area traits belong to the delayed group of displaced features. This group of displaced features is enriched in genes that denote high RNA metabolism. Features that may accelerate (early displacement) their expression pattern along the leaf development in the PK in comparison with C_3_ included VB diameter, OBS Area % and genes involved in catabolism, stress response and leaf development. Only three of a total of 15 phenotypic traits were conserved between C_3_ and PK species suggesting an important modification of the leaf anatomy at this stage of evolution.

While the number of features displaced from Proto-Kranz to C_4_ species shows similar trends to the displacement from C_3_ to Proto-Kranz, there is less difference between the number of these displacements. Most of the features displaced earlier in leaf development are enriched in genes involved in mRNA processing, among other functions. In contrast, features in the delay group are enriched in genes related to proteolysis, cell morphogenesis, and developmental cell growth. Interestingly, the transition from Proto-Kranz to C_4_ species suggests that most of the phenotypic traits are conserved, indicating that the leaf anatomy does not require significant adaptations to the C_4_ pathway.

### Transcription factors potentially involved in the regulation of IBS cell area

Addressing the displacement of leaf anatomical traits between species is another way of identifying potential regulators involved in particular pathways. We were interested in the displacement of the IBS area among species.

The area of the IBS is an interesting phenotypic trait postulated as one of the most important in Kranz anatomy, because it is where CO_2_ carboxylation takes place in C_4_ species (Christin *et al*., 2013, Lauterbach *et al*., 2019). Our results show that the phenotypic trait IBS cell area is preserved between C_3_ and PK species but it is displaced in C_4_ species. While the IBS cells reach their maximum area in S3 at the base of the leaf in C_3_ and PK species, they do the same in S5, closer at the tip of the leaf, in the C_4_. This delay of the expression correlated with the displacement of 41 genes involved in cell wall modification and pyruvate transport, among others. Indeed, two genes in this group are known to be part of suberin biosynthesis: an ortholog to AtFAR1/4 (fatty acid reductase) a member of alcohol-forming fatty acyl-CoA reductases (Domergue *et al*., 2010) and an ortholog to AtABCG16, a class of ABCG half-transporters that are required for the synthesis of an effective suberin barrier in roots and seed coats (Yadav *et al*., 2014). In addition, there are two transcription factors among the features displaced. One is a member of the Class 1 TCP transcription factor family (AtTCP7) that in *A. thaliana* plays an important role during leaf and hypocotyl development (Aguilar-Martinez *et al*, 2013). The second one (AtBFP4) is a member of the GeBP family and is involved in the defense response in *A. thaliana* (García-Cano *et al*., 2018). So far, the role of these two candidate genes on leaf development has not been explored in grasses. They seem to be good candidates for further experiments.

In conclusion, this is the first study showing how unsupervised machine learning can be used for the discovery of patterns of variations between phenotypic traits and transcription factors associated with the development and evolution of Kranz anatomy. Moreover, this methodology is fast, easy to implement and permits visualizing hidden patterns in complex data matrices. This method can be easily applied to other model and non-model species as well.

The use of SOM was useful to find candidate genes as potential enablers of Kranz traits differentiation. In addition, the methodology allowed us to detect displacements in gene regulatory networks between species with different photosynthesis subtypes. Studying the displacement of genes and phenotypic traits permitted us to identify drivers of phenotypic evolution. In fact, we identified transcription factors involved in the evolution of one of the most salient features of Kranz anatomy: the BS cell.

All genes proposed here might be good candidates for further validation by functional studies in grasses.

## Supplementary data

Table S1. Definition of quantitative phenotypic traits.

Table S2. Quality measures for SOMs

Fig. S1. Developmental gradient in the 5th leaf of four Otachyriinae subtribe species.

Fig. S2. Quantitative phenotypic traits

Fig. S3. SOM Interactive visualization graph

Fig. S4. SOM line visualization plot

Fig. S5. GO terms (BP) from Neurons 14 and 161 clustered by REVIGO.

Fig. S6. GO terms (BP) from Neurons 15 and 160 clustered by REVIGO.

Fig. S7. GO terms (BP) from Neurons 65 and 228 clustered by REVIGO.

Fig. S8. Species-specific SOM.

Fig. S9. Number of displacements.

Fig. S10. Phenotypic traits displacement.

Supplementary dataset 1. Phenotypic traits and gene expression data used for SOM analysis.

Supplementary dataset 2. Membership list of neuron pairs (SOM 17×17)

Supplementary dataset 3. GO enriched terms (BP) in C_3_ to PK and PK to C_4_ displaced features.

Supplementary dataset 4. Genes displaced together with IBS cell area and GO (BP) enriched terms.

## Abbreviations

BS: bundle sheath
GDC: Glycine decarboxylase complex
LGT: Lateral gene transfer
M: mesophyll
ML: machine learning
PK: Proto-Kranz
SOM: Self-Organizing Maps
VB: vascular bundle

## Acknowledgements

We thank members of the Development and Evolution lab (LED, IAL) and the Instituto de Agrobiotecnología del Litoral (UNL-CONICET) for helpful discussions. We also thank Juan Manuel Acosta for providing plant material. A special thanks to the FULBRIGHT program and the Ministry of Education of Argentina that made possible the realization of this work. We are grateful to anonymous reviewers for critically reading the manuscript.

## Author contributions

SP, AJS and RR defined the research question. SP, GS and RR contributed to the study design and experiments; GS helped with the SOM model design. SP performed the experiments; SP, AJS and RR contributed to the biological interpretation of the results; SP, GS and RR wrote the manuscript; SP, GS, AJS and RR revised the manuscript. RR is the corresponding author. All authors read and approved the final manuscript.

## Data availability

Data generated or analyzed in this study are included in this published article and its supplementary data published online.The Illumina RNA-Seq reads and transcriptomes assembly underlying this article are available in NCBI Sequence Read Archive (Prochetto *et al*., 2023; PRJNA813546).

